# The role of secondary pollinators in the evolution of complex colour signals in a bimodal pollination system

**DOI:** 10.1101/2025.06.01.657234

**Authors:** Katharine L. Khoury, Jon Ågren, Craig I. Peter, Nina Sletvold, Ethan L. Newman

## Abstract

Flower colour is recognized as a key trait influencing pollinator visitation and behaviour, evolved to match the sensory system of a particular pollinator group. However, some flowers combine colours associated with different pollinators, suggesting bimodal adaptation. The perennial herb *Cyrtanthus obliquus* produces red corollas with yellow tips and is pollinated by birds, but also by bees. To assess the role of the two pollinator categories in shaping selection on the bi-coloured corollas, we examined bee choices in arrays in which the colour signals of plants had been manipulated, and we quantified total as well as bird-mediated phenotypic selection on flower colour using both bird and bee visual models. Bees preferred flowers with a yellow signal over all-red flowers. The analysis of phenotypic selection indicated that both birds and bees influence net selection on flower colour, resulting in conflicting selection on the colour contrast between corolla and corolla tip. Our findings show that the optimal phenotype should depend on the relative importance of the two categories of pollinators in a given population, and are consistent with bimodal adaptation. This study fills an important gap by showing that secondary pollinators can contribute to selection on complex colour signals in angiosperm flowers.

## Introduction

Mounting evidence suggests that the diversification of floral traits within the angiosperms has been driven by pollinator-mediated selection [1, 2]. This is thought to have happened through divergent selection on floral traits due to differences in the choices and morphologies of different pollinators [3–6]. When flowers are visited by more than one pollinator, it is often assumed that the most frequent and efficient pollinators will exert the strongest selection and have the strongest influence on floral trait evolution [7–10], as stated in the most effective pollinator principle [7]. However, there are several examples of flowers with traits that are seemingly adapted to more than one pollinator and that do not fit a single pollination syndrome [11–14]. Among these are plants with bimodal pollination systems, i.e., plants pollinated by two different functional groups of pollinators [15–18]. In these plants, flowers may provide signals that are closely tied to the sensory system of more than one pollinator group, including scent [16], colour [19] or both [13, 15, 18, 20]. While such combinations of signals could represent adaptations to different pollinators, few have experimentally tested their functional significance, let alone, whether they are under selection by different pollinator groups.

Flower colour is a key trait defining pollination syndromes [21, 22], reflecting adaptation to the visual system of different pollinator groups. For example, bees lack the photoreceptors to detect long wavelengths of light in the red spectrum of flowers of bird-pollinated plants [23], whereas many nectarivorous birds are likely to perceive the entire spectrum between 300-700 nm [24]. For plants with bimodal pollination, differences in colour perception between pollinator groups implies that combinations of colour signals, that successfully attract both pollinator groups should be optimal. A complex colour signal is especially likely to evolve if resource costs of producing additional colours are weak, and if there are no fitness costs associated with the display of colours attractive to multiple pollinators (*sensu* [25]), i.e., if a secondary pollinator can be added without reducing the fitness gain by the primary pollinator. Such a scenario has been suggested to explain temporal colour shifts in flowers, where retaining old flowers of a new colour can attract additional pollinators [26, 27]. However, whether colour combinations displayed simultaneously represent adaptations to multiple pollinators remains largely unknown.

Linking observed selection to a specific pollinator group requires that other possible pollinators are experimentally excluded, which can be a major challenge in natural populations. To date, most studies attempting to separate the effects of different pollinators have taken advantage of systems with temporal separation in pollination, and isolated the contribution of diurnal and nocturnal pollinators by excluding them during night versus day [6, 28, 29]. Interestingly, these studies have documented that patterns of selection do not always follow expectations based on the MEPP. For example, although diurnal pollinators contributed most to seed production in the orchid *Gymnadenia conopsea*, diurnal and nocturnal pollinators mediated similar strength of selection on floral display and morphology, at least when considered in isolation [29]. At another site, pollinators mediated selection on both day- and night-emission rates of individual scent compounds, although, in this study, nocturnal pollinators contributed most to seed production [30]. This suggests that also less important pollinators contribute to selection. Nevertheless, few studies have isolated selection by distinct functional groups of pollinators in natural populations of any species, and it remains unknown whether secondary pollinators contribute to the evolution of ‘bimodal adaptations.

In this study, we examine the functional and adaptive significance of the combination of two colour signals in the giant fire lily, *Cyrtanthus obliquus* (Amaryllidaceae). There are reports that this species is pollinated by sunbirds [31], but also by pollen-collecting bees (pers. obs.). Its floral traits are primarily associated with the sunbird pollination syndrome, including long, red corolla tubes containing copious amounts of nectar [32]. However, unlike most other members of its genus that possess uniformly red corollas [33], the corolla of *C. obliquus* has yellow tips, which creates a notable contrast with the proximal, red corolla bases (Fig. 1). Yellow flower colour is often associated with adaptation to bees [34], and we hypothesize that the yellow corolla tips are important for attracting bees, as secondary pollinators of the fire lily. To test this idea, we combined two experimental approaches. First, we evaluated the importance of the yellow corolla tips for attraction of bees by removing or increasing the yellow signal. If the yellow tips have evolved to attract bees, removing the signal should reduce bee visits, whereas increasing the signal should increase bee visits. Second, we quantified selection on flower colour and morphology in an open-pollinated treatment where both bees and birds had access to flowers, and in a caged treatment, where birds were excluded. By comparing the two treatments, we obtained estimates of selection imposed by birds. In the open-pollinated treatment, where birds are expected to dominate among pollinators, we expected selection on the colour contrast between the red corolla and the background in bird vision, and for a corolla length corresponding to the bill length of sunbirds. In the caged treatment, where flowers are visited by bees only, we expected selection for a larger yellow corolla area, and increased contrast between the yellow corolla tip and the background, and between the red and yellow parts of the corolla in bee vision. Finally, we used supplemental hand-pollinations to test the hypothesis that pollen limitation is stronger in the caged than in the open-pollinated treatment, as expected if sunbirds are the main pollinators.

**Figure 1.**
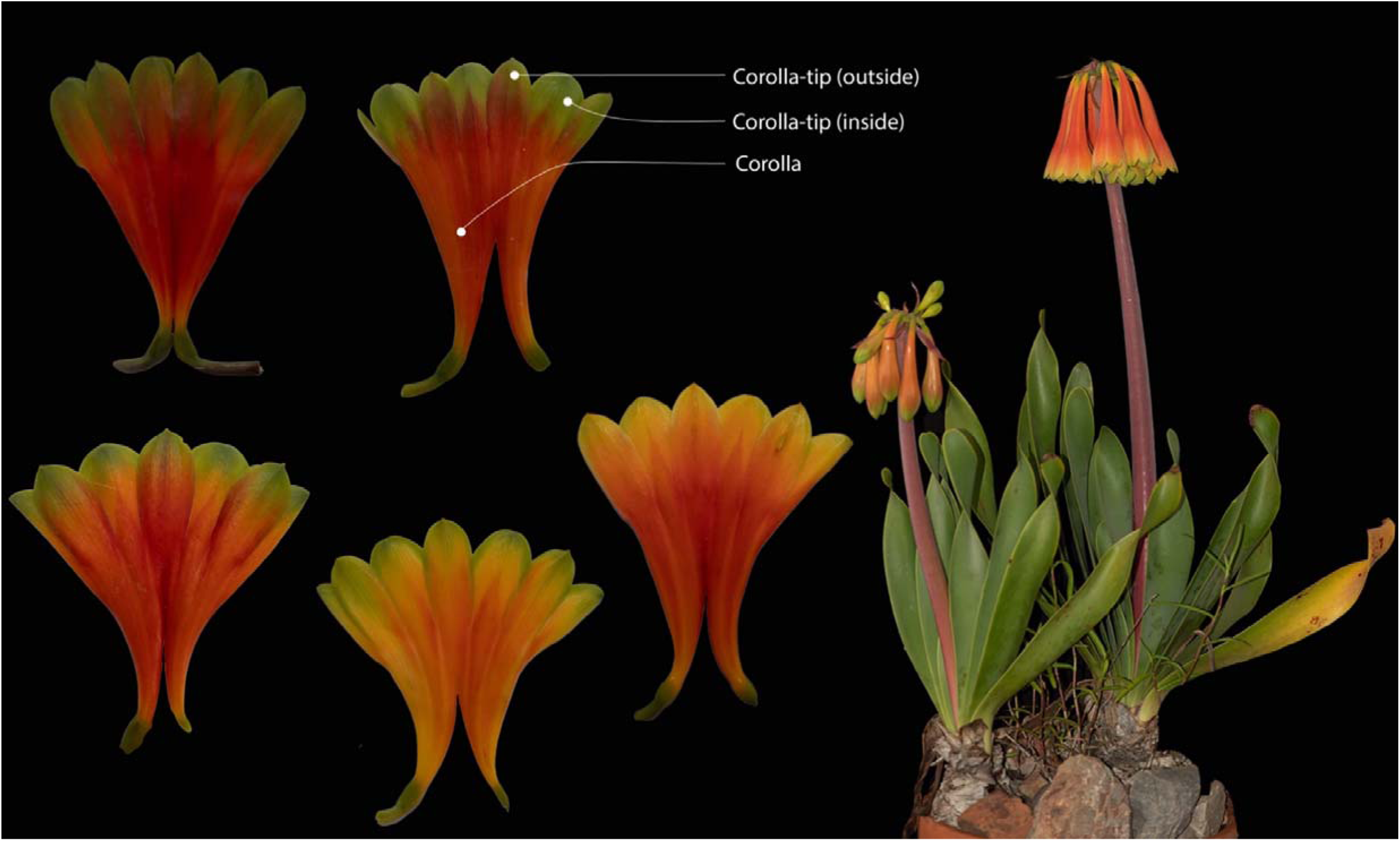
Colour plate illustrating the bicoloured flowers of *Cyrtanthus obliquus*, where the yellow-green tip contrasts with the red-orange corolla tube.

## Materials and Methods

### Study system

*Cyrtanthus obliquus* (L.f.) Aiton is a perennial bulbous geophyte that occurs in rocky outcrops in thickets and grasslands and is widespread in the Eastern Cape and southern KwaZulu-Natal Provinces of South Africa. Flowering occurs between October and December, and plants produce one to several inflorescences displaying multiple pendulous flowers with red to orange tubular corollas and prominent yellow-green tips (Fig. 1). The corolla consists of fused inner and outer tepals, and only the tips of the outer tepals are visible at the bud stage. In the study population, the ca 5 cm long corollas contain large amounts of nectar (mean ± SD, 78 ±58.01μL, n=241) with low sugar concentration (mean ± SD, 21.40 ±7%, n=240; measured using 10 μL glass capillary tubes (Hirschmann, Eberstadt, Germany) and an Atago 0–50% hand-held refractometer (Bellevue, USA; measurements conducted on one flower per sampled plant). The flowers have been suggested to be primarily adapted to pollination by malachite sunbirds (*Nectarinia famosa*) [35]. We conducted our study in a large natural population in Makhanda (−33.251156, 26.578503), where flowers are visited by both malachite sunbirds and bees. The sunbirds perch on the robust inflorescence stem while feeding on the nectar. Most visits from bees are made by the large carpenter bee (*Xylocopa sicheli*), that forage for pollen, although Lassioglossum and honeybees are sometimes observed. A breeding system experiment conducted in the study population showed that *C. obliquus* is self-incompatible (*see* Supplementary materials).

### Bee and bird perception of *C. obliquus* flower colour

To evaluate whether corolla colour might have been shaped by interactions with both birds and bees, we quantified how well birds and bees perceive the corolla and corolla tip against the background, and the corolla tip against the rest of the corolla, using data on spectral reflectance and models of bird and bee vision. We measured the spectral reflectance of the surface of the red part of the corolla, and of the yellow tips of the inner and outer tepals on 164 individuals. We used an Ocean Insight FLAME Miniature spectrometer (Ostfildern, Germany), with a PX-2 Pulsed Xenon Light Source and a Premium 400μm fibre optic probe. We also took three spectral measurements of the surrounding vegetation and two measurements of rocks from the study site, of which the average was used as an estimate of background reflectance in the colour vision models. The raw reflectance spectra for different parts of the flower were plotted using the “*aggplot*” function in the R package ‘*pavo’* [36].

Colour vision models were then used to determine how birds and bees perceive the corolla colours of *C. obliquus*. Both the tetrahedral colour space [37] and the receptor noise limited model [38] are commonly used as visual models for birds [39], while for bees the colour hexagon [40] is the preferred visual model. For visual modelling in both bee and birds, we used both the colour spaces and the receptor noise limited model [41, 42]. All spectral processing, visualisation and colour vision modelling were completed in the R package ‘*pavo’* [36]. All data analyses for this study were conducted in R v.4.1.2 (R Core Team, 2024).

#### Bee perception

Bees have a conserved trichromatic visual system [43], with UV, blue and green photoreceptors [23]. We modelled bee colour perception using the *vismodel* function with D65 daylight illumination and imported background spectra to calculate hyperbolically transformed quantum catches for each photoreceptor. Reflectance spectra were plotted in the trichromatic colour hexagon for the *Apis mellifera* visual system [44] using the *colspace* function. We then calculated Euclidean distances between points within the hexagonal colour space using the *coldis* function. Spectra that fall into different quadrants of the Chittka [44] trichromatic colour hexagon are considered distinguishable by bees. However, bees have continuous colour discrimination, and they can reliably detect differences between colours within the same quadrant for distances larger than 0.11 hexagon units [45]. In addition to calculating Euclidean distances, we used receptor noise limited (RNL) models to calculate noise-weighted colour distances for the *Apis* visual system [38]. The quantum catches were used to calculate the noise-weighted colour distances in Just Noticeable Differences (JNDs) using the *coldis* function. JNDs larger than 0.27 are considered to be distinguishable by bees [46].

#### Bird perception

Birds have a tetrachromatic visual system, with UV, blue, green and red photoreceptors [47].We obtained quantum catches for each photoreceptor using the same import variables as for the bee model. Quantum catches were plotted in tetrahedral bird colour space using the *colspace* and *tcsplot* functions. Similar to bees, the *coldis* function was used to calculate Euclidean distances between points within bird tetrahedral colour space. As for bees, we additionally used the RNL model to calculate noise-weighted colour distances [38]. For the RNL models, as for the tetrahedral colour space, we used the *coldis* function to calculate noise-weighted colour distances (in Just Noticeable Differences, JNDs). Colour distances larger than 1 JND were assumed to be distinguishable by birds [38].

For both models, the colour distance between the tip and the rest of the corolla (Euclidean distances and JNDs) was calculated based on measurements taken on a single individual.

### Bee flower colour preference

To determine if the yellow corolla tips of the fire lily increase attractiveness to pollen-collecting bees, we set up experimental arrays in the natural population. In this experiment, we manipulated flower colour using paints (Dala craft paint) that matched the natural red and yellow corolla colours (assessed by spectral reflectance measurements, see Supplementary material). We included a total of four colour treatments: uniform yellow corolla (increasing the yellow signal), uniform red corolla (removing the yellow signal), red corolla with yellow tips (painted control), and unpainted control. Each array included six inflorescences arranged in three treatment pairs. Within an array, pairs were separated by one meter, and inflorescences within a treatment pair by 20 cm. In the treatment pairs, visitation to the painted control and to one of the three other colour treatments was compared, resulting in three binary choices: 1) the painted control against the uniform red corolla, expecting a preference for the painted control, 2) the painted control against the uniform yellow corolla, expecting a preference for the uniform yellow, and 3) the painted control against an unmanipulated corolla (true control), expecting no difference in preference. The third pairing was included to test the effect of paint *per se*. Furthermore, the painted control does not seem to be associated with an increased painted contrast in comparison to the naturally occurring contrast in the bee colour hexagon (*P* > 0.05, Fig. S1, Methods S1). Significance values associated with pairwise contrasts are reported in Fig. S1.

To set up the choice experiment, we collected at least 50 fresh inflorescences in the study population, each displaying at least three open flowers containing pollen, and we used all inflorescences within three days. All inflorescences were trimmed to six flowers to control for effects of resource availability on pollinator choices. In the painted treatments, all flowers in the inflorescence were painted the evening before we set up arrays in the field. After colour manipulations, each inflorescence was mounted upright in a plastic jar containing water. We applied paint to all jars, which were hidden to an approaching pollinator, to control for potential effects of paint fragrance on bee visitation. Arrays were set up in the morning, and observed between 08:00 and 14:00 on days with temperatures above 25°C. An array was arranged in a crescent around two observers, with one person taking notes while the other observed pollinator choices. When a foraging pollinator entered the array, we recorded the bee species, its identity, and the visitation sequence. We defined the first flower visited within any colour treatment pair as a first choice. We regularly swopped the positions of treatments within and between pairs to prevent bias in choice based on the direction in which bees approached the array. After several visits by pollen-collecting bees, an inflorescence pair was replaced within the array. We conducted the arrays during peak flowering, over a total period of five days in November 2021 and 2022. At each date, we used new inflorescences that had been painted the night before.

Bee colour preferences were analysed using generalized linear mixed effects model with a binomial error distribution and a logit link function, using the *glmmTMB* function, implemented in the ‘glmmTMB’ package [48]. We analysed each binary choice (treatment pair) separately. In these models, pollinator identity was nested within year as a random effect, and first choices were used as the response. In all models, we used the *Anova* function from the ‘car’ package [49] to test for significance of model terms and we used the *emmeans* function from the package ‘emmeans’ [50] to obtain back-transformed confidence intervals, which we plotted using ‘ggplot2’ [51]. All models were checked for overdispersion using the “simulateresiduals” function using the ‘DHARMa’ package [52], which revealed no concerning trends in the model residuals.

### Context-dependent selection: separating the contribution from birds

#### Experimental design

To examine whether selection on floral traits differed between plants exposed only to bees, versus to both birds and bees, we conducted a selective pollinator exclusion experiment between October and December 2022. We marked a total of 207 flowering plants that all had a single inflorescence (plants with more than one inflorescence per bulb were rare in the natural population), and randomly allocated individuals to a bird-exclusion treatment (n = 96) and an open-pollinated control treatment (n = 111). In the exclusion treatment, plants were individually covered with wire cages (2.5 by 2.5 cm mesh) throughout the flowering season. For each plant, we recorded the number of flowers, inflorescence height, and the length of the corolla tube. The latter was measured from the base of the ovary to where the corolla flares, to the nearest millimetre using digital callipers (Fig. 1). In addition, we sampled one flower per plant for measurements of colour traits. First, we took spectral measurements of the adaxial sections of the red and yellow parts of the corolla and determined the contrast between the red corolla and the background in bird vision, the contrast between the yellow corolla tip and the background in bee vision, and the contrast between the yellow tip and the red part of the corolla in both bird and bee vision, as described above. The choice of contrast traits was based on the results of the initial models of bee and bird perception (see below).

We used the hexagon and tetrahedral visual models for bees and birds, respectively. Second, we dissected and flattened the flower underneath a thin pane of glass (Fig. 1), and photographed it using a DSLR (Canon 5D MK4 with a 100mm macro lens). The scale was determined with a steel ruler placed under the glass pane. Images were processed in ImageJ [53], and we used the polygon tool to calculate the area of the yellow tips as well as the total corolla area. We quantified yellow signal strength as the proportion of the corolla that was yellow. At the end of the flowering season, we collected all fruits produced, dissected ripe fruits in the lab using a surgical blade and counted all seeds. We used total seed production per plant as a measure of female fitness. Some plants missed records of phenotypic traits or fitness, and the final sample size was 84 in the caged treatment and 72 in the control treatment. We quantified pairwise correlations between all eight measured traits in each treatment using the Pearson correlation coefficient. To limit the number of model terms and avoid problems of collinearity, we excluded three traits that were strongly correlated to another trait in one or both treatments, i.e., plant height (height vs. number of flowers; r_open_=0.47, r_caged_=0.62), contrast between red corolla and yellow tip in bird vision (contrast corolla and tip vs. contrast corolla and background; r_open_=0.81, r_caged_=0.71) and contrast between the yellow corolla tip and the background in bee vision (contrast tip and background vs. contrast corolla and tip; r_open_=0.79, r_caged_=0.60).

#### Phenotypic selection

We quantified selection on the five floral traits (number of flowers: 9.36±2.85, 30.4%; corolla tube length:49.35 ±4.77, 9.67%; yellow corolla proportion: 22.53±3.66, 16.25%; tip-corolla contrast in bee vision: 0.140±0.076, 54.99%, corolla-background contrast in bird vision: 0.34±0.11, 32.35% - mean±SD, CV) using the multiple regression approach of Lande and Arnold [54], with relative fitness (individual number of seeds produced divided by mean seed production) as the response variable, and standardized traits (with a mean of zero and variance of one) as explanatory variables. Relative fitness and standardized traits were calculated separately by treatment. We estimated directional selection gradients β*_i_* from models including only linear terms, and quadratic selection gradients γ_ii_ from models including both linear and quadratic terms. For quadratic selection gradients, we doubled the coefficients extracted from the regression model [55]. We visualized the form of selection using partial regression plots. To examine if selection in the two pollination treatments differed, we used an ANCOVA model including relative fitness as the response variable and the five standardised traits and trait by pollination treatment interactions as explanatory variables. Bird-mediated selection was quantified as Δβ_bird_ =β_open_ - β_caged_, where β_open_ is the selection gradient of open-pollinated plants visited by both birds and bees, and β_caged_ the selection gradient of plants visited by bees only. Due to a large number of zeroes in the dataset, we used nonparametric bootstrapping to estimate 95% confidence intervals of β_open_, β_caged_, and Δβ_bird_ by the bias-corrected and accelerated (BCa) method [56]. We generated 10 000 bootstrap estimates for each treatment combination. Confidence intervals for estimates of bird-mediated selection (Δβ_bird_) were obtained by calculating pairwise differences between the 10 000 bootstrap estimates of β_open_ and β_caged_. In all cases, we based statistical significance on CI not overlapping zero.

#### Pollen limitation

To quantify pollen limitation and determine whether pollen limitation increased when birds were excluded, we compared seed production of supplementally hand-pollinated plants with seed production of plants in the two pollination treatments in the selection study. In the supplemental hand-pollination treatment, we included 30 individuals at the start of their flowering, and saturated the stigmas of all open flowers with pollen from at least five individuals that were located more than 10 meters away from the focal plant. Hand-pollinations were repeated every two days until all flowers on the 30 individuals had received pollen. We recorded fruit set and harvested fruits at maturation, and determined number of seeds per individual by dissecting fruits and counting seeds in the lab. In both pollination treatments, we calculated pollen limitation as 1-(mean seed production in open or caged pollination treatment/mean seed production in the hand-pollination treatment). We used a generalized linear model with a Poisson error distribution and a log link function to examine if seed production significantly differed between the three pollination treatments using the stats package. A *Posthoc* tukey test was conducted in the package “emmeans”, and plots were generated in “ggplot2”.

## Results

### Bee and bird perception of complex colour signals

Averaged reflectance spectra varied among flower parts, and the results of the two visual models (hexagon and tetrahedral vs. RNL) were very similar, for in both bee and bird vision. The red outer surface of the corolla had a UV absorbent curve that peaked around 650nm (Fig. S2). This contrasted strongly with the yellow tips of the inner and outer tepals that peaked at around 550nm, including a minor UV peak in measurements of the inner tepal (Fig. S2). In bee vision, the most discriminable contrasts were the outer corolla tip against the red corolla and the outer corolla tip against the background, in both visual models (Fig. 2A,B).

**Figure 2.**
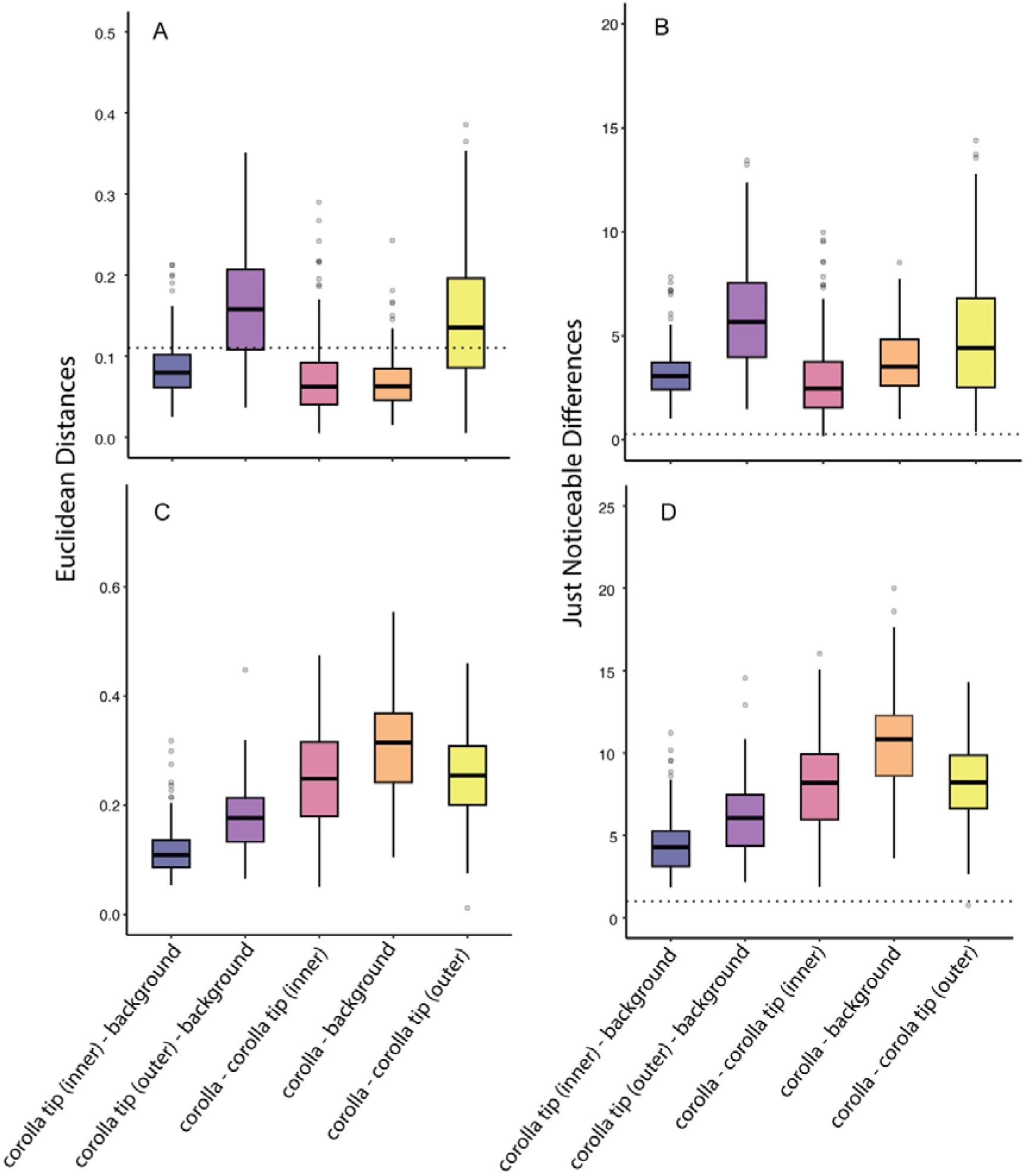
Colour contrasts between *Cyrtanthus obliquus* flowers and the background and between different parts of the flower likely involved in attracting pollinators. Colour contrasts are measured as A) Euclidean distances in hexagonal bee vision, B) just noticeable differences in the receptor noise model associated with bee perception, C) Euclidean distances in tetrahedral UV bird vision, D) just noticeable differences in the receptor noise model associated with bird perception. Dotted lines in each panel represents the perceptibility threshold for each of the colour vision models, with values above the threshold considered to be perceptible.

According to the hexagon model, the red corolla against the background, the inner corolla tip against the background, and the inner corolla tip against the red corolla were all poorly distinguished by bees (Fig. 2A). For the receptor noise models, all JND values were above 0.27, indicating that all calculated colour distances were perceptible by bees (Fig. 2B). All colours were perceptible in bird vision, using both the tetrahedral model with UV, and the receptor noise-limited model, where all contrasts were above the JND perceptual threshold of one (Fig. 2C, D). The most discriminable contrasts were the red corolla against the background and both the inner and outer corolla tips against the red corolla (Fig. 2C, D).

### Bee flower colour preference

In total, 48 *X. sicheli* bees and solitary bee (31 individuals in 2021 and 16 in 2022) made a total of 163 first choices in the binary choice experiment. Bees significantly preferred the painted control over the uniform red corolla (χ^2^ =10.51, df =1, *P =*0.0012; Fig. 3A), and made 35 first choices in favour of the painted controls, compared to 18 first choices of the red corollas. Bees also tended to prefer the uniform yellow flowers over the painted control (χ^2^ =3.2, df =1, *P =*0.07; Fig. 3B), with 43 first choices of yellow flowers and 32 of painted controls. Finally, bees tended to prefer painted controls over true controls (χ^2^ =2.76, df =1, *P =*0.097; Fig. 3C), with 21 first choices by bees of painted controls compared to 14 first choices of unmanipulated plants. This suggests an effect of paint *per se*, that likely was due to increased total reflectance of painted compared to natural flowers (Fig. S2).

**Figure 3.**
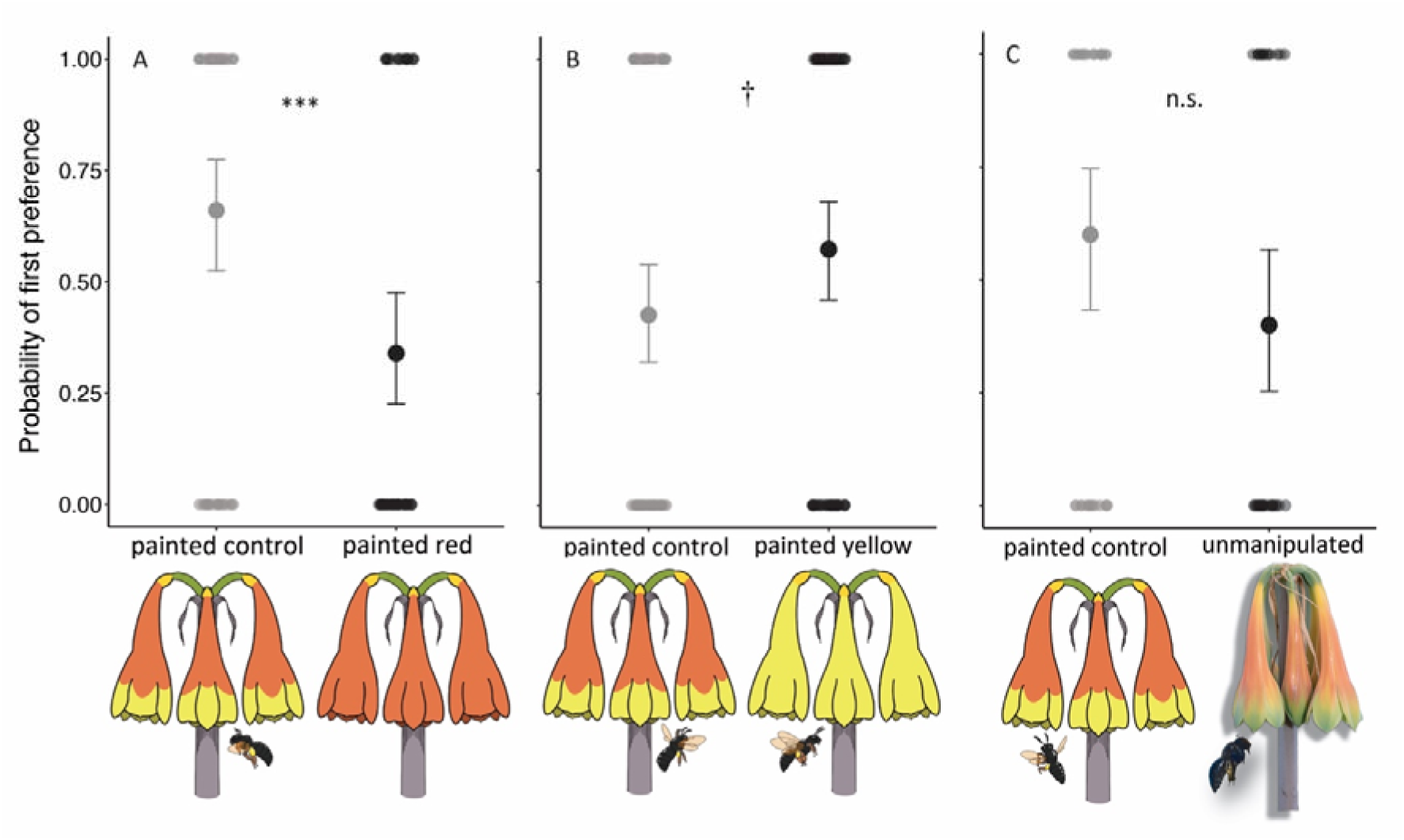
Bee preferences for flower colour of *Cyrtanthus obliquus* in the binary choice arrays with painted flowers: A) uniform red flowers compared to painted controls (bicoloured flower), B) uniform yellow flowers compared to painted controls, C) painted control compared to natural flower (bicolured flower without paint).

### Pollen limitation and phenotypic selection

We found strong pollen limitation of seed production in both the open-pollinated and caged treatments, and pollen limitation was significantly stronger when birds were excluded (PL_open_ = 0.65, PL_caged_ = 0.93; χ^2^ =106.37, df = 3, *P<*0.0001, Fig.S4).

Among plants available to both bird and bee pollinators, there was directional selection for more flowers (β*_open_* = 0.28±0.12, Table 1), and stabilizing selection on colour contrast between the tip and the corolla in bee vision, with an optimum close to the current mean (γ*_open_* = −0.37±0.19, Fig. 4). There was no significant linear or quadratic selection on any other trait (Table 1). Among plants available to bees only, we detected directional selection for increased colour contrast between the tip and the corolla in bee perception (β*_caged_* = 0.34±0.19, Fig. 4). There was a statistically significant negative quadratic estimate of selection on number of flowers (Table 1), and nonlinearity was due to a decelerating increase in fitness with higher number of flowers (no intermediate optimum).

**Figure 4.**
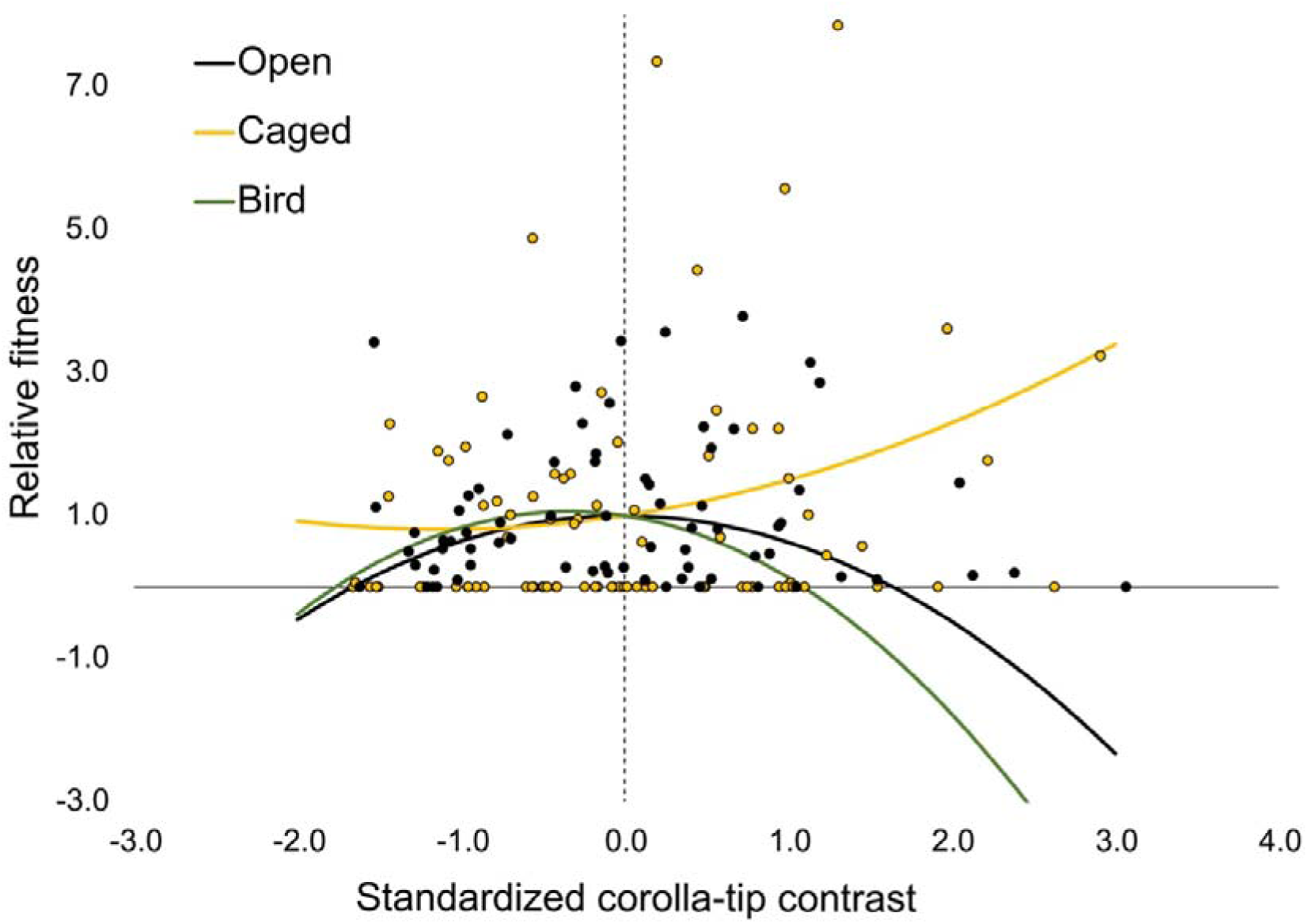
Selection on colour contrast between the red corolla and the yellow corolla tip among open-pollinated plants (net selection, black line and dots), and among caged plants where birds were excluded (yellow line and dots). The resulting bird-mediated selection calculated as the difference between treatments is also illustrated (green line). Associated statistical tests are given in Table 1.

**Table 1.** Phenotypic linear (β*_i_*±SE) and quadratic (γ*_ii_*±SE) selection gradients on floral traits (number of flowers, corolla length, proportion of yellow on corolla, contrast red corolla vs. yellow tip in bee vision, contrast red corolla vs. background in bird vision) of *Cyrtanthus obliquus* with 95% bootstrapped CI among plants exposed to an open-pollinated treatment visited by both birds and bees, and a caged treatment where birds were excluded. Bird-mediated linear (Δβ*_i-bird_*) and quadratic (Δγ*_ii-bird_*) selection gradients were calculated as the difference between treatments. Significant selection gradients (95% CI not overlapping zero) are indicated in bold.

| Trait | Open $\beta_i \pm SE$ | Low<br>CI | High<br>CI | Open $\gamma_{ii} \pm SE$ | Low<br>CI | High<br>CI | Caged $\beta \pm SE$ | Low<br>CI | High<br>CI | Caged $\gamma_{ii} \pm SE$ | Low<br>CI | High<br>CI | $\Delta\beta_{i-bird}$ | Low<br>CI | High<br>CI | $\Delta\gamma_{ii-bird}$ | Low<br>CI | High<br>CI |
| --- | --- | --- | --- | --- | --- | --- | --- | --- | --- | --- | --- | --- | --- | --- | --- | --- | --- | --- |
| No. Flow | **0.281 ± 0.119** | 0.011 | 0.46 | -0.102 ± 0.167 | -0.58 | 0.22 | 0.223 ± 0.174 | -0.03 | 0.53 | **-0.398 ± 0.316** | -0.89 | -0.01 | 0.058 | -0.58 | 0.74 | 0.295 | -0.34 | 0.82 |
| Corolla | 0.132 ± 0.119 | -0.09 | 0.34 | 0.093 ± 0.170 | -0.62 | 0.12 | 0.236 ± 0.186 | -0.07 | 0.54 | -0.219 ± 0.276 | -0.72 | 0.24 | -0.104 | -0.98 | 0.52 | 0.312 | -0.30 | 0.94 |
| Pr. yellow | 0.013 ± 0.121 | -0.23 | 0.24 | 0.127 ± 0.205 | -0.22 | 0.50 | 0.054 ± 0.204 | -0.33 | -0.45 | 0.493 ± 0.326 | -0.28 | 1.20 | -0.041 | -1.02 | 0.78 | -0.367 | -1.16 | 0.48 |
| Tip-Cor | -0.007 ± 0.123 | -0.22 | 0.20 | **-0.367 ± 0.185** | -0.68 | -0.06 | **0.342 ± 0.187** | 0.07 | 0.65 | 0.152 ± 0.271 | -0.34 | 0.60 | **-0.350** | -1.45 | -0.02 | -0.519 | -1.08 | 0.04 |
| Cor-Back | -0.137 ± 0.119 | -0.35 | 0.08 | 0.040 ± 0.240 | -0.56 | 0.44 | 0.103 ± 0.181 | -0.16 | 0.37 | -0.046 ± 0.303 | -0.54 | 0.52 | -0.240 | -1.16 | 0.20 | 0.086 | -0.72 | 0.70 |

Comparisons of the two treatments showed that birds mediated directional selection for reduced colour contrast between the tip and the corolla in bee vision (Δβ*_bird_* = −0.35), in combination with stabilizing selection for an optimum that is lower than the current mean (Δγ*_bird_* = −0.52; Fig. 4). However, estimates of bird-mediated selection were not statistically significant (Table 1).

## Discussion

Bimodal pollination systems have been suggested in several species based on observed effective pollination by two functional groups and floral traits apparently adapted to pollination by each of them [15]. Here, we experimentally demonstrated that the yellow corolla tips of the bird-pollinated fire lily *Cyrtanthus obliquus* attract bees that act as secondary pollinators, and that both functional groups influence net selection on flower colour. Conflicting selection on flower colour indicates that there is an adaptive tradeoff between becoming successfully pollinated by birds and bees, and that the optimal phenotype should depend on the relative importance of these two categories of pollinators in a given population.

By producing floral trait combinations that attract flower visitors that differ in morphology, behaviour and sensory system, plants may maintain sufficient pollen transfer in environments where the composition of the pollinator community varies in space and time [57–59]. Still, it has been suggested that such trait combinations should be transitional, because adaptive tradeoffs should lead to specialization on the most important pollinator [7, 10]. In the fire lily population, the malachite sunbird is clearly the most important pollinator, as documented by the strong increase in pollen limitation when birds were excluded (Fig. S4). So, what maintains the bicoloured corollas of *C. obliquus*? Estimates of directional phenotypic selection in the presence and in the absence of bird pollinators showed that corolla-tip colour contrast in bee vision was subject to conflicting selection. As expected, bees, possibly together with other environmental factors, mediated selection for increased corolla-tip contrast. However, unexpectedly, birds mediated selection for reduced contrast in combination with stabilising selection for an optimum below the current population mean (Fig. 4). Why birds mediate selection on a trait modelled in bee vision is unclear. One possibility is that the higher pollination intensity in the open-pollination treatment leads to expression of resource costs associated with the production of two colours, which could explain the observed selection for reduced contrast. Alternatively, the selection is due to correlations with traits not included in the analysis. To examine this possibility, we calculated pairwise correlations between corolla-tip contrast in bee vision and colour traits in bird vision. Corolla-tip contrast in bee vision was positively correlated with tip-background contrast in bird vision (r=0.56), suggesting that a certain corolla-tip contrast in bee vision may actually be favoured by the birds because of selection on this correlated trait. If so, the tradeoff between attraction of birds and bees is not as strong as it would otherwise be. Instead, the bird-mediated stabilising selection should contribute to the maintenance of the bicoloured corollas and an increased chance of attracting bees as secondary pollinators, since the preference experiments indicated that the yellow corolla tip of the red flower is important for the attraction of bees. In the open-pollinated treatment, where both birds and bees visited the flowers of the fire lily, the conflicting linear selection cancelled out, and only stabilising selection on corolla-tip contrast remained. This suggests that both birds and bees contribute to maintaining this trait close to its current mean.

The theoretical framework for when specialisation in pollination should be favoured is based on pollinator-mediated tradeoffs [60, 61], where models in their simplest form assume that effects of multiple pollinators on strength and direction of selection are additive, and are weighted by their fitness contributions. Our estimate of bird-mediated selection (selection in open-pollinated treatment minus selection in bird-exclusion treatment) also assumes additivity. The estimate of bird-mediated stabilizing selection on corolla-tip contrast was very similar to that documented in the presence of both pollinators (Fig. 4), which is in line with expectations of parallel contributions to selection and fitness, and the most important pollinator principle [7]. However, interactive effects of pollinators on fitness and selection can be expected whenever removal of pollen by one species reduces that of another, and have been predicted to be strongest when an efficient pollinator interacts with one that mainly acts as a pollen thief (the ‘good’ vs. ‘ugly’ pollinator [62]). Moreover, previous studies that separated selection by nocturnal and diurnal pollinators in more generalized systems did not find strictly additive effects of the two pollinator categories [28, 29], indicating the presence of interactions. To test rigorously whether effects on selection by birds and bees are additive in *C. obliquus*, also the effect of bees on selection would need to be experimentally isolated.

Besides corolla-tip contrast, we did not detect any evidence of bee- or bird-mediated selection on flower colour. In the choice experiment, bees preferred flowers that were painted completely yellow compared to painted bi-coloured controls, consistent with an innate preference for yellow [34]. However, although there was selection for increased corolla-tip contrast, there was no statistically significant selection on the proportion of total corolla area that was yellow, when only bees were visiting flowers. This suggests that the 2.3-fold variation in proportion yellow in the population was not enough to affect the likelihood of bee-mediated pollination success, and that the presence of yellow was sufficient for attraction of bees. Neither was there any evidence of bird-mediated selection on colour contrast between the red part of the corolla and the background, or on corolla tube length, despite that these traits are expected to influence bird attraction and morphological fit to the flower. These results suggest that all flowers in the population are highly visible to birds against the background, and that the efficiency of pollen transfer is similar across the range of flower depths present in the population. In many bird-pollinated species, corolla length is correlated with the beak length of the main bird pollinators [63–65]. However, in *Costus spiralis*, there was no difference in seed production among flowers visited by three hummingbirds with different beak length [66]. To clarify if current trait means are close to the optimum for bee and bird pollination success, an experimental increase in trait variation could be helpful [5, 59, 67].

Within the context of pollinator-mediated speciation, intermediate traits may act as an adaptive bridge to facilitate pollinator shifts[59]. Stebbins [7] referred to this process as the “intermediate step of double function” in pollinator-mediated speciation. The premise of this idea is that it would be easier to incrementally shift from one pollinator to another by evolving intermediate traits, instead of leaping between two distinct adaptive optima separated by a fitness trough. Flower color shifts are likely to involve strong tradeoffs, as documented in *Mimulus*, where colour had contrasting effects on visitation rates of bees and hummingbirds [68, 69]. Bicoloured corollas may represent such an intermediate trait, which reduces the strength of adaptive tradeoffs. It is possible that within bicoloured populations where birds become less abundant or absent, selection will favour increased attraction of bees and the evolution of pure yellow flowers. However, in the present study, we did not detect any bee-mediated selection on the proportion of the corolla that was yellow. This suggests that a transition from bicoloured to yellow flowers could be the result of the absence of bird-mediated selection favouring a red signal, rather than selection by bees.

In the genus *Cyrtanthus*, many species have completely red flowers and are likely pollinated by sunbirds [33]. However, the genus also includes several species with uniformly yellow or cream flowers visited by lepidopterans and bees, and mapping of flower colour onto the phylogeny suggests several independent colour shifts from red to yellow or cream [33]. Furthermore, there are at least two other species that resemble *C. obliquus* in having a red corolla with contrasting yellow-green tips, namely *C. herrei* and *C. tuckii* [31]. The latter species occurs in a pure red form that is likely visited by birds only, and a red form with green and yellow tips, which may attract both birds and bees. Phylogenetic analyses suggest that these three species are distantly related, and belong to separate clades in the genus [33]. The current phylogeny is poorly resolved, and further work is required to determine whether bicoloured corollas are likely to represent stable bimodal adaptations subject to stabilising selection or transitional states.

Only a few studies have quantified selection on colour traits using visual models of the primary pollinators [70, 71]. By using chromatic colour contrasts as traits, and by selecting traits based on their visibility in the perceiver’s sensory system, we should maximise the ability to detect pollinator-mediated selection on colour [72]. The accuracy of colour vision models can depend on variation in illumination and background colour [73, 74]. In particular, the contrast between flower and background colours can affect the perceptibility of colour traits [45]. Here, we used the actual background colours from the natural population in our visual models, which should improve the characterisation of colours as perceived by the pollinators, compared to using a standard green background.

By combining a detailed characterization of flower colour with two experimental approaches, we have provided several lines of evidence supporting a bimodal pollination system in *C. obliquus*, where both birds and bees contribute to pollen transfer and net selection on flower colour. This may represent a stable bimodal system, where the importance of bees will vary with bird pollination intensity, and where the bees, which are less-effective pollinators, provide a safeguard against complete pollination failure in the absence of birds.

## Acknowledgements

We are grateful to Tony Dold for introducing us to the study locality and to Anina Coetzee for helping us with visual modelling.

## Funding

This project was supported by a Thuthuka grant# 138321, a Rhodes University Research Council grant, and a STINT initiation grant# IB2023-9249.

## Statement of authorship

All authors conceived and planned the experiments and data collection. E.N. and K.K. carried out the experiments and data collection, K.K., N.S., and E.N. performed statistical analyses. EN wrote the first draft with input by KK on the methods. All authors contributed to conceptualizing the study, writing, editing and finalizing the paper.

## Notes

### Competing Interest Statement

The authors have declared no competing interest.

