## Supplementary Materials for "The role of secondary pollinators in the evolution of complex colour signals in a bimodal pollination system"

**Supplementary Methods and Figures**

**Methods**

**Breeding system**

To determine the breeding system of *C. obliquus*, we compared the supplemental hand-pollination treatment (outcrossing) in the main experiment to a controlled self-pollination treatment. Before flowering started, we bagged nine individuals with fine mesh. As flowers opened and stigmas became receptive, we saturated stigmas with self-pollen every other day, and continued the treatment until all flowers had wilted. All inflorescences remained bagged until fruit maturation in the population. None of the self-pollinated flowers developed a fruit (Fig. S4), indicating that *C. obliquus* is self-incompatible.

**Colour match between paints and natural flowers**

To ensure our painted treatments matched the natural flower colours in the *C.obliquus* study population, we compared the spectra of the applied paints (Dala craft paint) to the naturally occurring flowers in bee perception. We painted fresh flowers from eight individuals with both red and yellow paint from the corolla to the lowest tip of the tepals. The red paint was selected to match the corolla of *C. obliquus*, and the yellow paint was selected to match the corolla tips. On each individual, we took four spectral reflectance measurements of the flowers: the painted red on top of the red corolla, the painted red on top of the yellow tip, the painted yellow on top of the red corolla, and the painted yellow on top of the yellow tip. Colours did not differ when painted on a red versus yellow flower parts, and we only present red on top of red and yellow on top of yellow. All measurements were cleaned, processed and plotted in bee colour vision using the same procedure as in our array experiment. To compare the contrast between corolla and tip in painted and natural flowers, we calculated the Euclidean distances between the red and yellow parts of the corolla for each individual (Fig. S1). We used a generalized linear model with a gamma distribution to test for a difference between the painted and unpainted contrast in bee vision.

The spectral reflectance curves of the painted red corolla and the painted yellow corolla tip, largely match the reflectance curves of the natural corolla and outer corolla tip, respectively (Figure S2). However, the paints have a higher total reflectance than the naturally occurring corolla and outer corolla tip. When the paints and the naturally occurring flowers were modelled in bee colour vision, there were significant differences between the painted and unpainted contrasts within bee vision (*𝛘^2^* = 269.95, *df* = 5, *P* < 0.0001). Specifically, there were significant differences between the unmanipulated and painted contrast between the corolla and background (*P* < 0.0001, Figure S1). However, there were no significant differences between the unmanipulated and painted contrast between the corolla and the corolla tip (*P* > 0.05, Figure S1), or between the corolla tip and the background (*P* > 0.05, Figure S1).

**Figures**

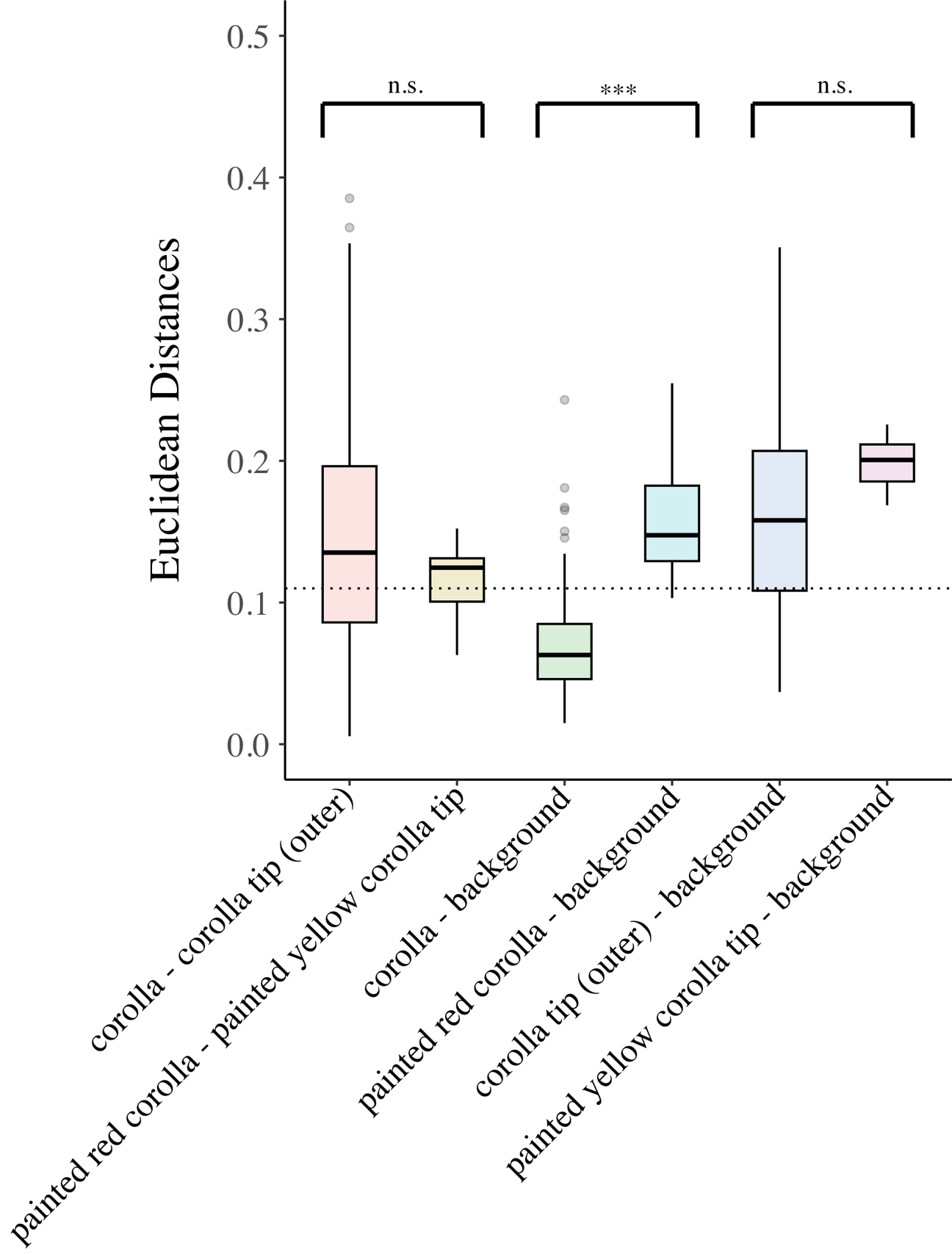

**Figure S1.** Euclidean distances in bee colour vision between the red corolla and the yellow outer corolla tips of natural, unmanipulated flowers and of painted flowers of *Cyrtanthus obliquus*. The perceptual threshold is 0.11 hexagon units, indicated by the stippled line.

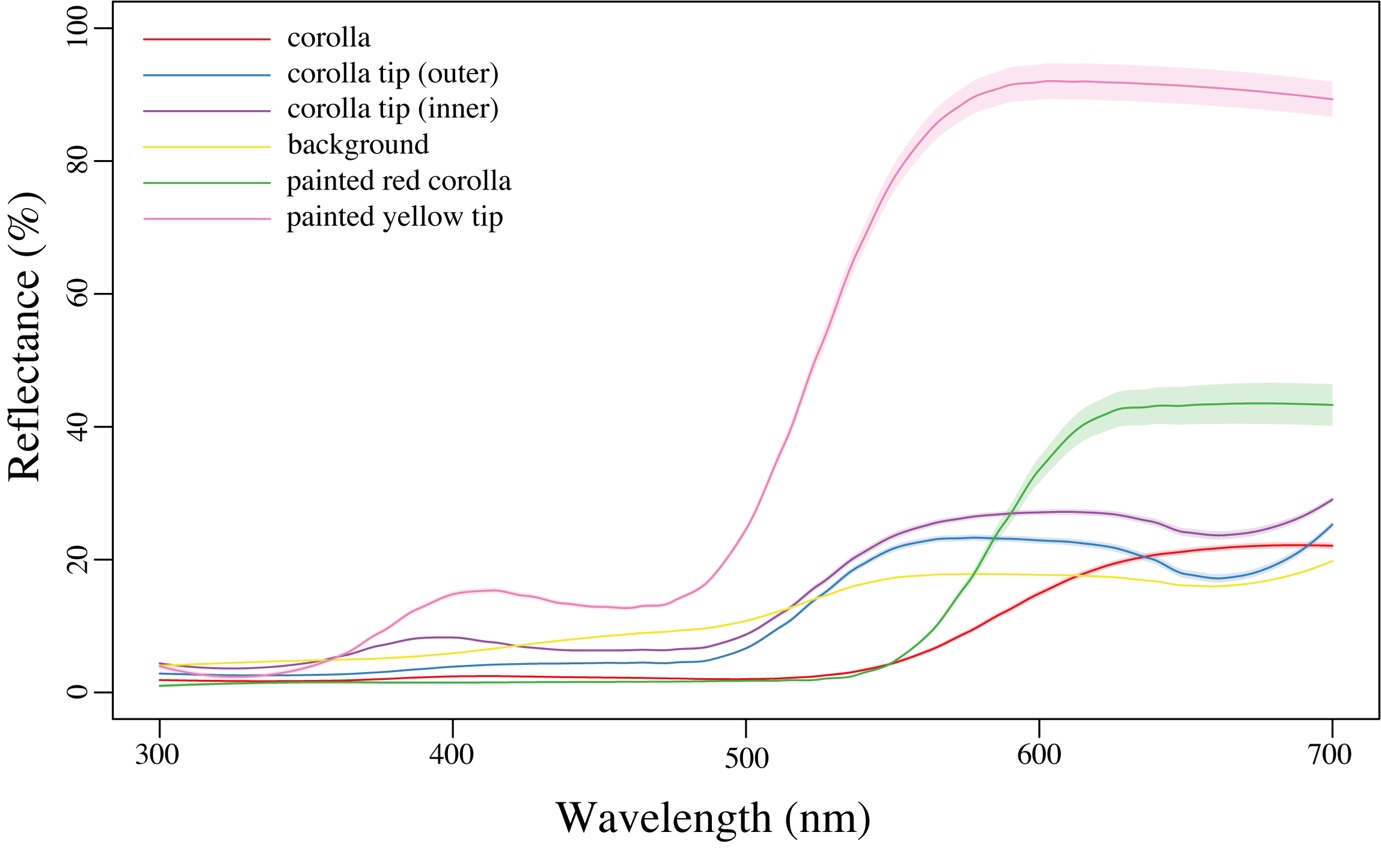

**Figure S2.** Spectral reflectance curves showing the mean and standard error of all measured surfaces on *Cyrtanthus obliquus* flowers and of the habitat background. The *C. obliquus* surfaces includes both unmanipulated and unpainted flowers.

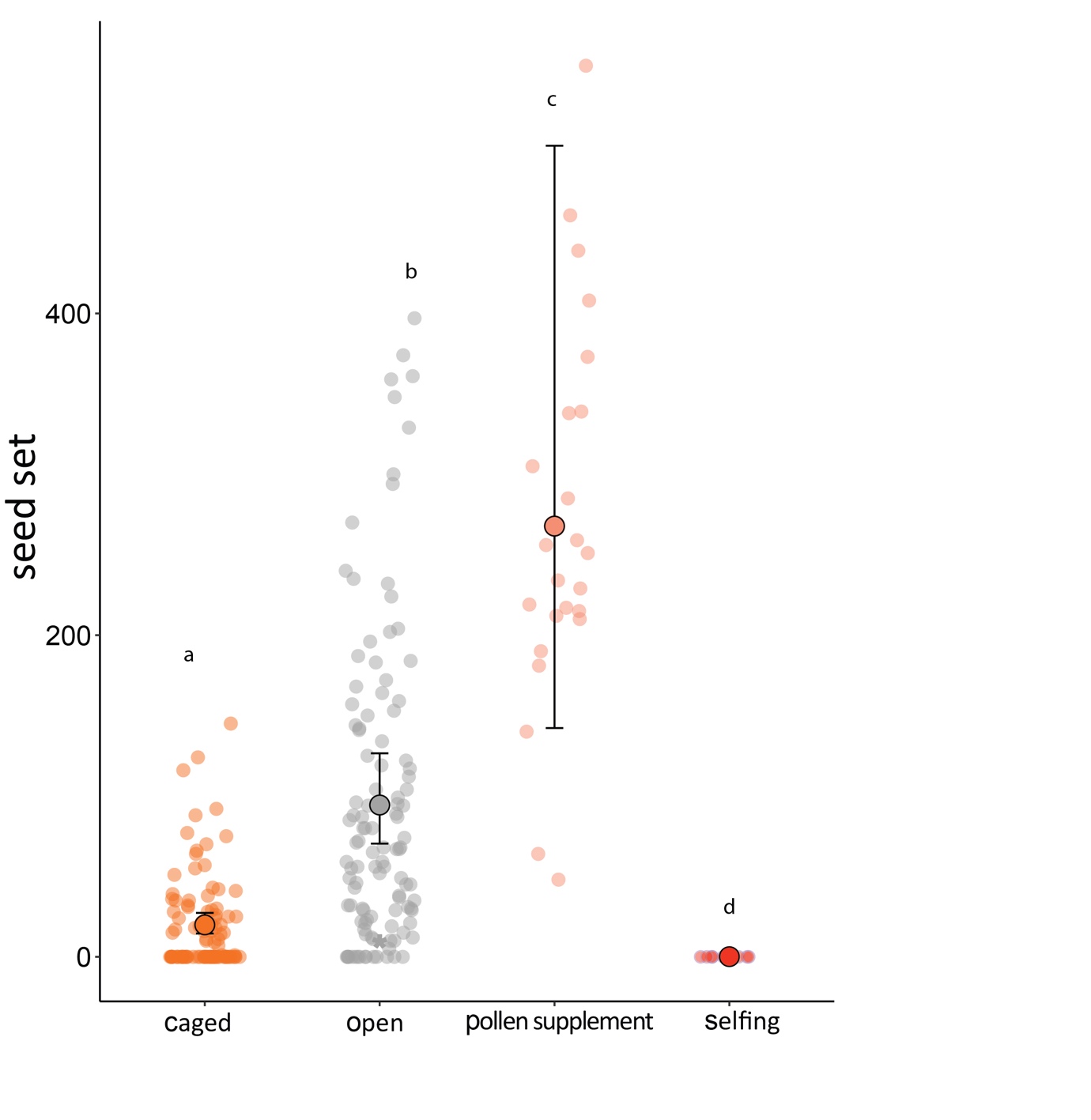

**Figure S4.** Seed set (total number of seeds) among caged plants (n=97), open-pollinated plants (n=112), supplementally hand-pollinated plants (n=24) and self-pollinated plants (n=9). Large circles represent mean seed set with asymmetric error bars, backtransformed from logit scale. Small, faded circles represent individual data points.
